## Supplementary Information for "Estimating effects of parents’ cognitive and non-cognitive skills on offspring education using polygenic scores"

### Supplementary Notes & Figures

**Deviation from pre-registered methods** 2

**Meta-analyses of Cognitive Performance and Educational Attainment GWAS** 3

**Supplementary Figure 1. GWAS-by-subtraction Genomic-SEM model, reproduced from Demange et al. 2020** 4

**Comparison of demographic and early-life characteristics of the adopted and non-adopted samples** 5

**Supplementary Figure 2. Birthplace coordinates for adoptees and non-adopted individuals.** 7

**Supplementary Figure 3. All direct and indirect genetic effect estimates** 8

**Supplementary Figure 4. Population genetic effects** 9

**Supplementary Figure 5. Ratios indirect effect/population effect** 10

**Supplementary Figure 6. Effect of the NonCog and Cog polygenic score on educational outcomes in monozygotic and dizygotic twins in TEDS and NTR** 11

**Supplementary Figure 7. Estimated effect of NonCog and Cog PGS on Educational Attainment in UK Biobank depending on the number of siblings (adopted or full-siblings) of the individual.** 12

**Methods for simulating genetic and phenotypic data in the presence of different biases and components** 13

**Comparison of two implementations of the sibling design using simulation** 14

**Supplementary Figure 8. Estimation of the direct and parental indirect genetic effects in simulated data with different implementations of the sibling design.** 16

**Comparison of sibling, adoption, and non-transmitted allele designs in presence of simulated components and biases** 17

**Supplementary Figure 9. Simulation results: Comparison of sibling, adoption, and non-transmitted allele designs in presence of components and biases** 19

#### Deviation from pre-registered methods

To correct for family structure in our trio data in NTR, we planned to use the gee function in R. To correct for additional shared factors in the sibling design, we planned to use the lme function in R to specify a random intercept for family (as done by (Selzam et al. 2019)). However, the gee function led to convergence issues when bootstrapping. Additionally, simulations showed that the use of a mixed model (lme or lmer commands in R) in the sibling design leads to underestimation of indirect genetic effects, and underestimation of the direct genetic effects in the non-transmitted alleles design. Hence, we use linear models in the sibling and non-transmitted allele design (lm command in R), and we bootstrapped standard errors (see Methods in the main manuscript).

#### Meta-analyses of Cognitive Performance and Educational Attainment GWAS

Before performing GWAS-by-subtraction, we ran a GWAS of Educational Attainment and Cognitive Performance in UKBiobank (polygenic score sample left-out). Genetic associations were obtained using fastGWA and controlling for age (Data-Field 21022), sex, array and the 25 first principal component analyses, for 1,246,531 SNPs (Hapmap 3 SNPs).

We then meta-analysed our UKB EA GWAS (N=388,196) with the EA GWAS by Lee et al excluding 23andMe, UK Biobank and NTR cohorts (N=318,916) using the METAL software. We included SNPs with sample-size > 500,000 and MAF > 0.005. We did not apply genomic control, but following Lee et al. we inflated the standard errors from the meta-analysis by the square root of the LD score intercept (1.223, SE=0.0223). After inflation of the SEs, we found a SNP heritability of 0.1006 (SE=0.0027) with a LD score intercept of 0.9783 (SE=0.0187).

We meta-analysed our UKB CP GWAS (N=202,815) with CP GWAS by Trampush et al (N=35,298) using the METAL software. We included SNPs with sample-size > 100,000 and MAF > 0.005. We found a SNP heritability of 0.1858 (SE=0.0064) with LDSC. We did not apply genomic control, and because the LD score intercept was acceptable (1.055, SE=0.0118), we did not inflate standard errors.

#### Supplementary Figure 1. GWAS-by-subtraction Genomic-SEM model, reproduced from Demange et al. 2020

Procedure identical to Demange 2020, description repeated verbatim for convenience: Cholesky model as fitted in Genomic SEM, with path estimates for a single SNP included as illustration. SNP, Cognitive performance (CP) and Educational attainment (EA) are observed variables based on GWAS summary statistics. The genetic covariance between CP and EA is estimated based on GWAS summary statistics for CP and EA. The model is fitted to a 3x3 observed variance-covariance matrix (i.e. SNP, CP, EA). Cog and Non-Cog are latent (unobserved) variables. The covariances between CP and EA and between Cog and NonCog are fixed to 0. The variance of the SNP is fixed to the value of 2pq (p = reference allele frequency, q = alternative allele frequency, based on 1000 Genomes phase 3). The residual variances of CP and EA are fixed to 0, so that all variance is explained by the latent factors. The variances of the latent factors are fixed to 1. The latent variables were then regressed on each SNP that met QC criteria.

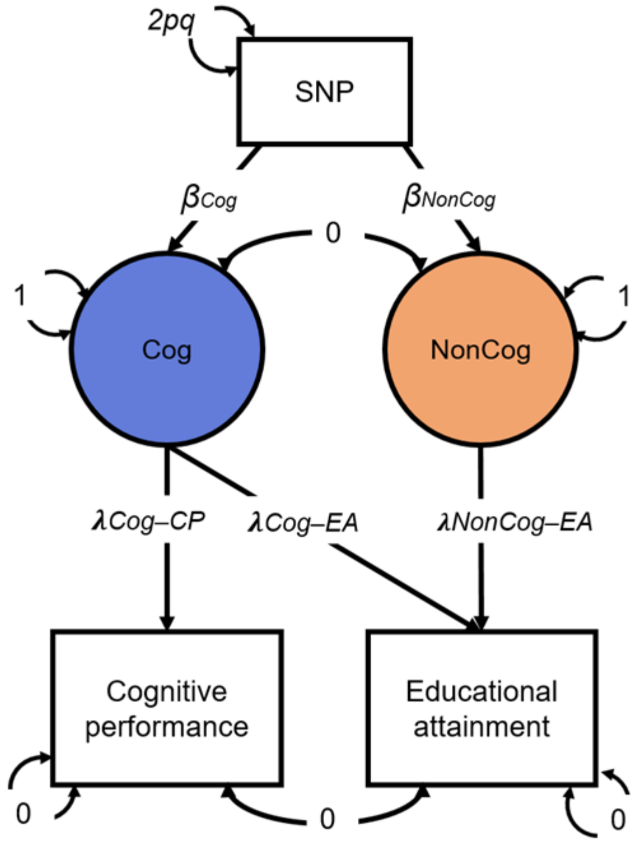

#### Comparison of demographic and early-life characteristics of the adopted and non-adopted samples

In the adoption design, indirect genetic effects are inferred by subtracting polygenic score associations estimated in a sample of adoptees away from those estimated in a non-adopted control group. When taking the difference, it is important that the groups are similar in characteristics other than genetic relatedness to their parents. We explored this empirically by comparing demographic and early-life characteristics of adoptees and nonadopted individuals in the UK Biobank.

**Supplementary Table 11A** displays results from our comparison of *NonCog* PGS, *Cog* PGS, birthweight, and educational attainment. We observed significant but small differences between the groups in their mean *NonCog* and *Cog* PGS as well as educational attainment (Cohen's d < 0.15). We do not observe differences in the variances of these measures between the two groups.

We observed differences in birthweight, with adopted individuals being lighter at birth than the non-adopted individuals (mean = 3.12kg vs 3.33kg, cohen's d=0.31). The variance in birthweight was also significantly different, with more variance in the adoptees group (0.57 vs 0.45). However, missing data is high in both groups, and in particular in adoptees (72% of missing data among adoptees, 43% in the non-adopted group).

We investigated whether the birthplaces of adoptees and non-adoptees were clustered differently. If so, this could mean that population stratification effects are not consistent across the groups. Hence, we performed k-means clustering on the UK Biobank's east/west birthplace coordinates, separately for adoptees and non-adopted individuals. Without a hypothesis for the number of geographical clusters, we used different numbers of clusters (k), and then plotted the within-cluster sum of squares according to k. For both adopted and non-adopted groups, the best number of clusters was 4, indicated by the location of the bend in the plot. As shown in **Supplementary Figure 2 (below)**, the pattern of clustering was also the same between the groups, with the clusters reflecting the regions of Wales, the South, the North, and the Midlands.

Next, we compared the proportions of adopted and non-adopted individuals born in each UK region (see **Supplementary Table 11B**). Adoptees were less likely to have been born in the Midlands and more likely to have been born in the South, but these differences were small. Since there could be differences between the groups in location of birth *within* each region, we also compared the cluster means for the two groups. There were only small differences. The largest discrepancy was that adoptees from the north were born further south than the non-adopted northerners (northern coordinate 1.93 vs 1.99).

Notably, the interpretability of these analyses are hindered by data limitations. A large fraction of birthweight data was missing (72% missingness for the adoptee group). Also, birthplace was self-reported, so could be inaccurate, particularly for adoptees.

Overall, the key variables under study are well-matched between the adopted and non-adopted groups. There may be systematic differences relating to birthweight and geography, but it is difficult to see how these would influence our results. We therefore are tentatively confident that taking the difference in the estimates between adopted and non-adopted groups is a valid approach.

#### Supplementary Figure 2. Birthplace coordinates for adoptees and non-adopted individuals.

Note: adopted (Left) and non-adopted (Right); colours = regions obtained by k-means clustering of coordinates.

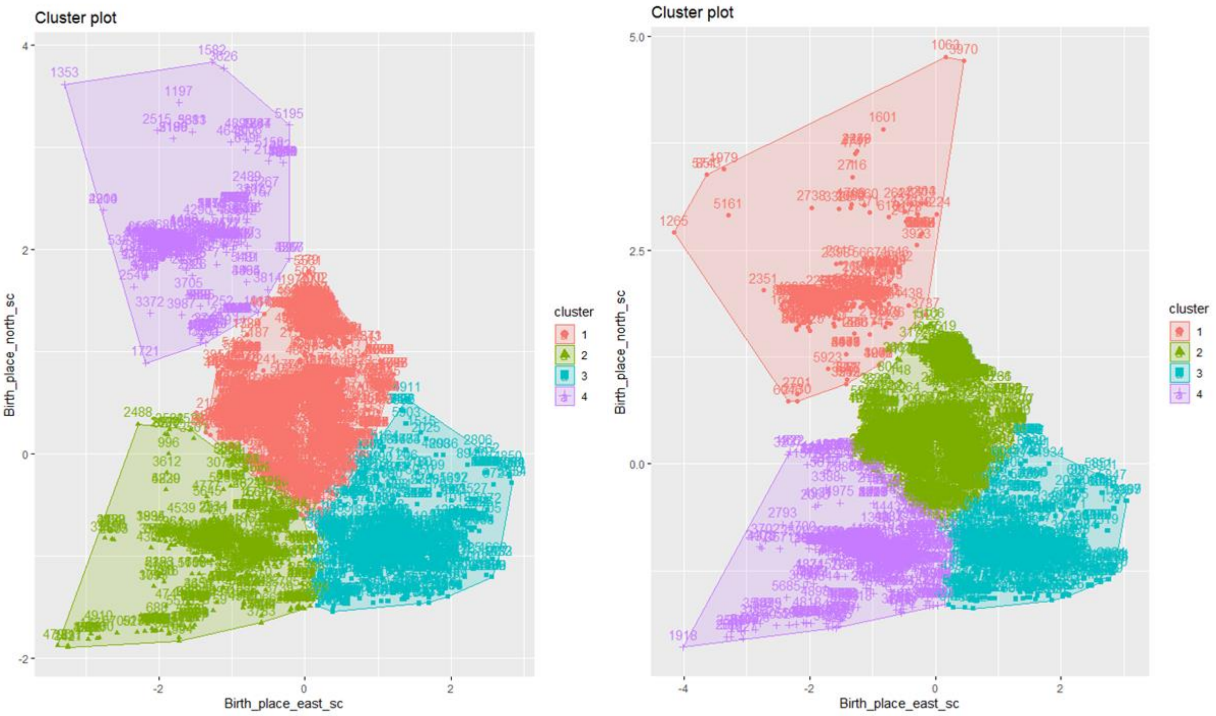

#### Supplementary Figure 3. All direct and indirect genetic effect estimates

Estimates of direct and indirect effects of *NonCog* and *Cog* PGS for every condition. Method is represented by a symbol, sample and outcome are represented by a color, bars are 95% CIs. Precise values are presented in **Supplementary Table 3.**

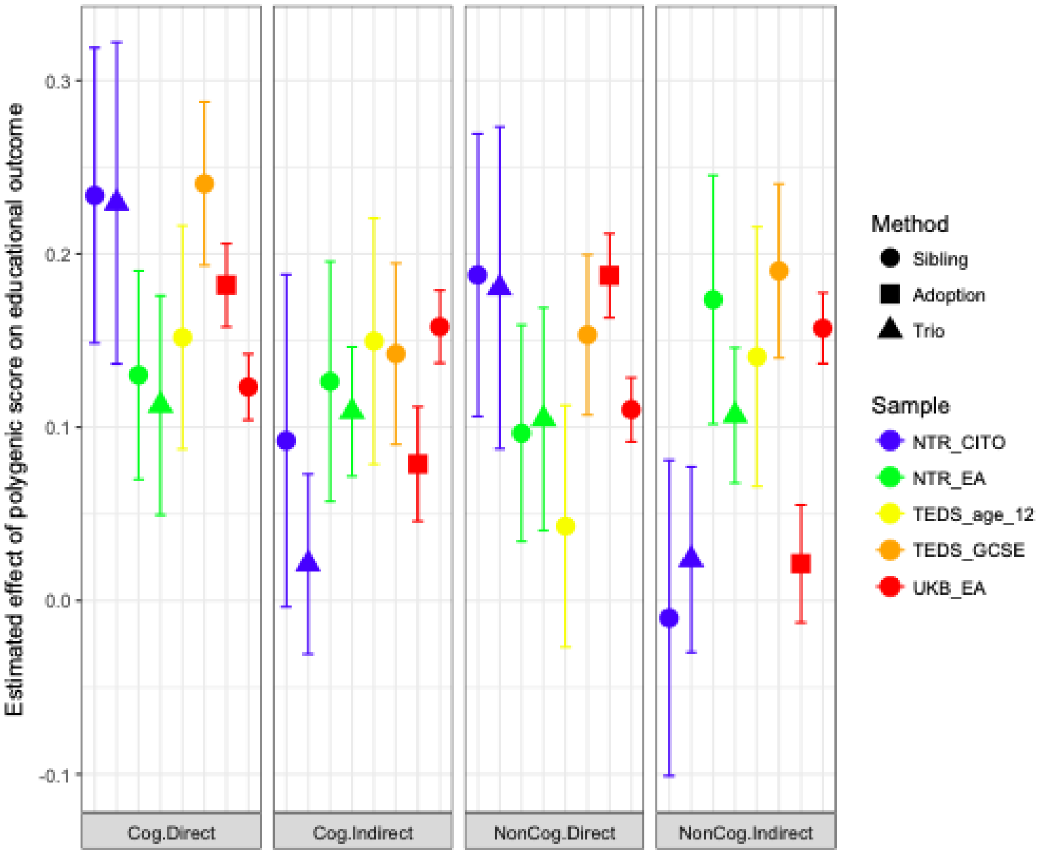

#### Supplementary Figure 4. Population genetic effects

Estimates of population effects of *NonCog* (orange) and *Cog* (blue) PGS for every condition grouped by method. X axis ticks indicate the sample (NTR, TEDS and UKB) and outcomes (CITO is age 12 achievement in NTR, 12yo is age 12 teacher rated achievement in TEDS, GCSE is age 16 achievement in TEDS and EA), bars are 95% CIs. Values are in **Supplementary Table 3.**

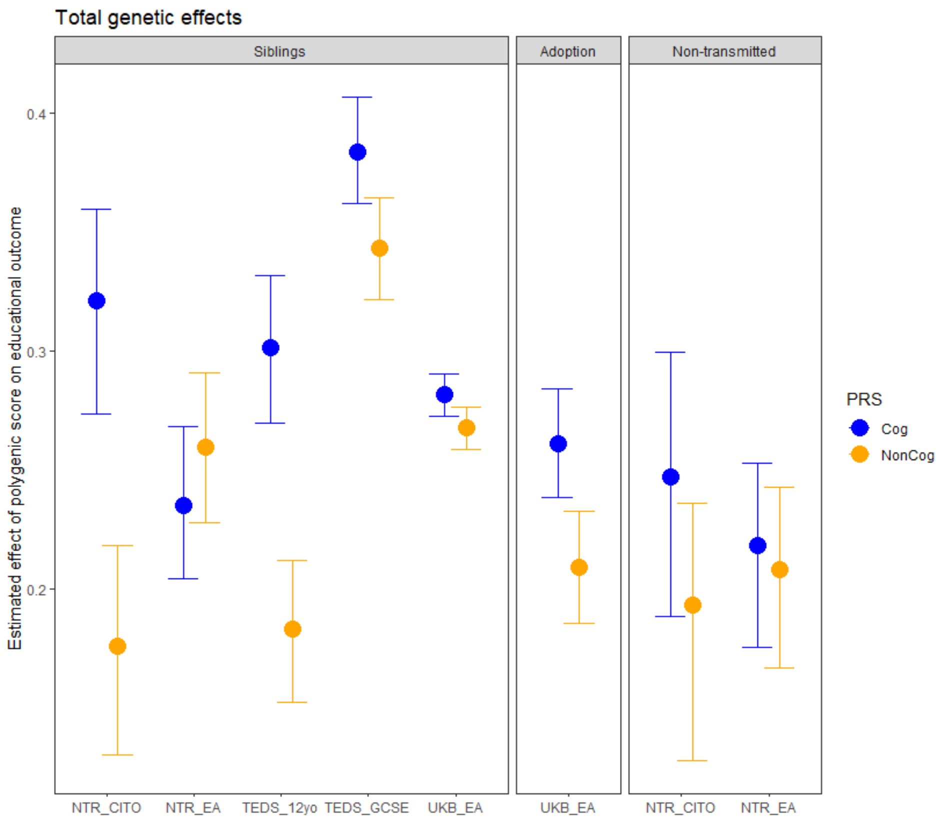

#### Supplementary Figure 5. Ratios indirect effect/population effect

Estimates of the ratio of the indirect effects on the population effects of *NonCog* (orange) and *Cog* (blue) PGS for every condition grouped by method. X axis ticks indicate the sample (NTR, TEDS and UKB) and outcomes (CITO is age 12 achievement in NTR, 12yo is age 12 teacher rated achievement in TEDS, GCSE is age 16 achievement in TEDS and EA); bars are 95% CIs. Values are in **Supplementary Table 3.**

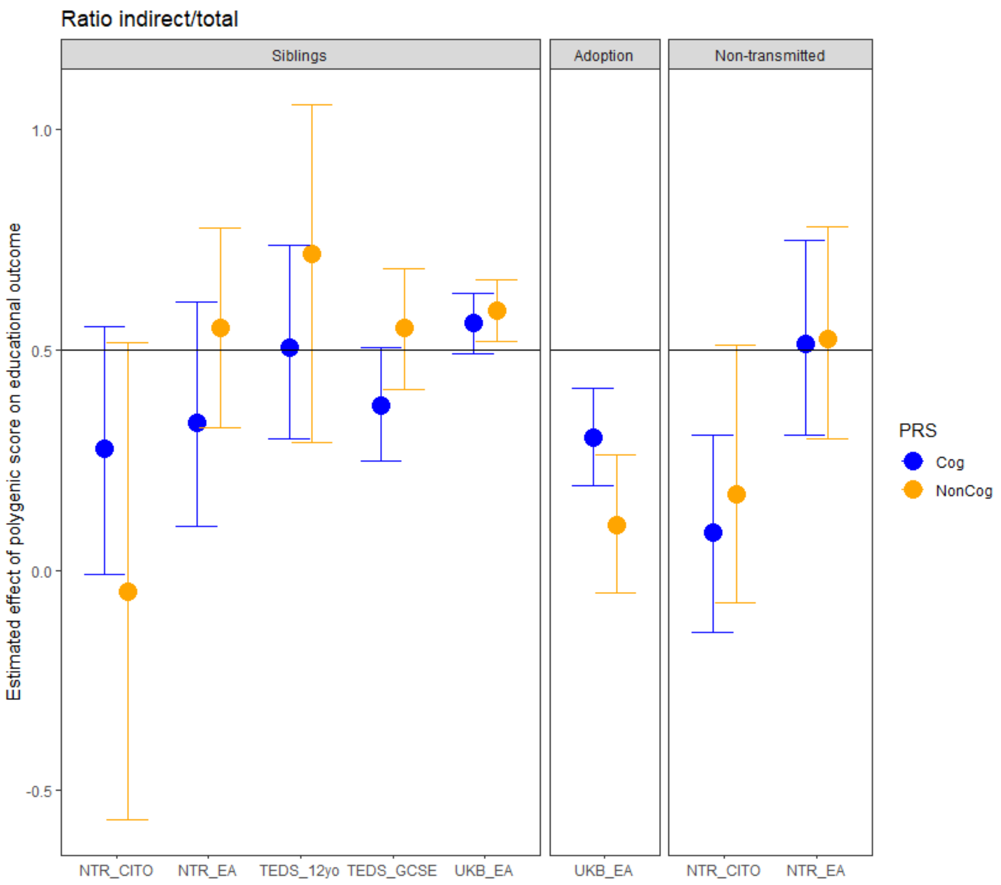

#### Supplementary Figure 6. Effect of the *NonCog* and *Cog* polygenic score on educational outcomes in monozygotic and dizygotic twins in TEDS and NTR

**A. Effect of *NonCog* PGS on educational outcomes in MZ vs DZ.**
**B. Effect of *Cog* PGS on educational outcomes in MZ vs DZ.**
Y axis represents the beta coefficient of the regression of the *NonCog/Cog* PGS on educational outcomes. In blue are results in TEDS and green in NTR. Bars represent the standard errors of the estimate. X axis ticks indicate the individual zygosity (MZ: monozygotic twin; DZ: dizygotic twin) and outcomes (age12 is age 12 teacher rated achievement in TEDS, age16 is age 16 achievement in TEDS, EA is educational attainment in NTR and CITO is age 12 achievement in NTR)

A.

B.

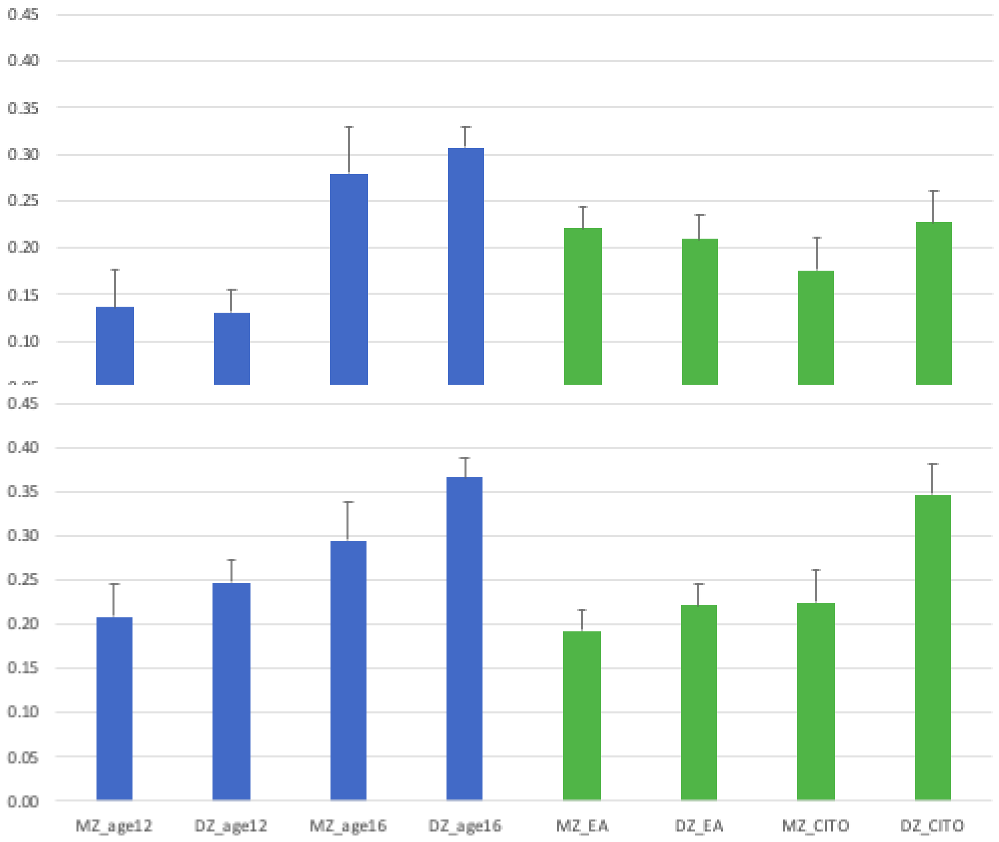

#### Supplementary Figure 7. Estimated effect of NonCog and Cog PGS on Educational Attainment in UK Biobank depending on the number of siblings (adopted or full-siblings) of the individual.

We look at the effect of *NonCog* and *Cog* PGS in non-adopted and adopted individuals (the latter group providing a control scenario since they are not genetically related to their sibling(s)). Blue dots represent the estimates of the *Cog* polygenic score effect (Beta) on educational attainment, bars are 95% CI. In orange are *NonCog* estimates. We draw the linear regression model between these estimates (blue and orange lines).

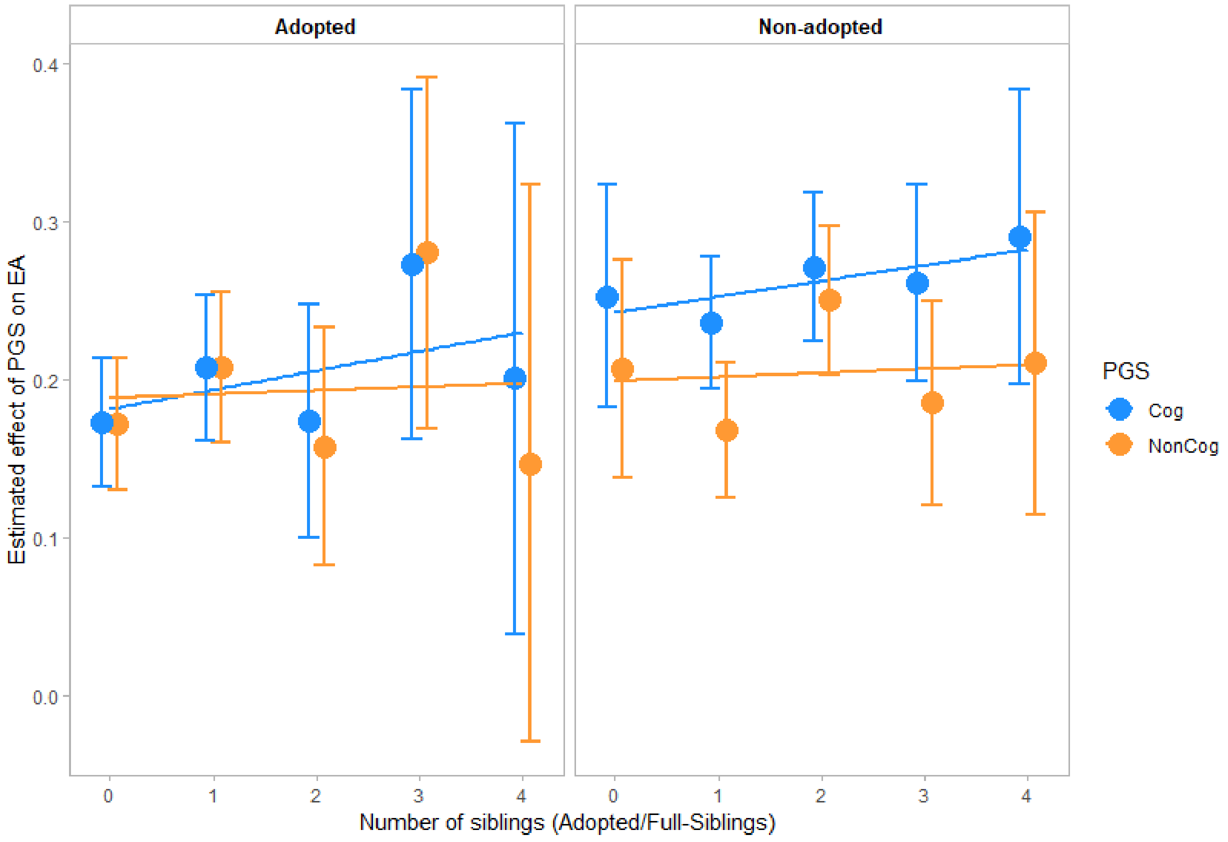

#### Methods for simulating genetic and phenotypic data in the presence of different biases and components

We simulate data introducing various potential (environmentally mediated) genetic effects and confounding biases, we then fit all models used throughout the paper in order to identify how the estimated parental indirect effect changes in the presence of these biases. We simulate 100 bi-allelic SNP calls for a sample size of 20,000 families. Each family includes a mother, a father, a biological child, a child sibling, and an adopted child sibling. The adoptee genotypes are drawn from another simulated dataset of biological parents, independent of the focal families. We create polygenic scores for simulated individuals by weighting genotypes by GWAS effects, which are defined as true SNP effects sizes plus error. **We simulate nine offspring traits influenced by different factors:**

i) direct genetic effects only,
ii) direct and indirect parental genetic effects (maternal and paternal),
iii) indirect parental genetic effects plus a prenatal indirect maternal genetic effect,
iv) indirect sibling genetic effect,
v) indirect parental genetic effects and an indirect sibling genetic effect,
vi) assortative mating,
viI) assortative mating and an indirect parental genetic effects,
viii) population stratification,
ix) population stratification and indirect parental genetic effects.

Having simulated the traits as detailed below, we then use three designs (sibling, adoption, non-transmitted allele; as detailed in the main article) to estimate indirect parental genetic effects on each trait. This allows us to evaluate how designs are affected by the components (prenatal and postnatal parental indirect genetic effects) and biases (sibling indirect genetic effects, assortative mating, population stratification). We repeated the simulation 100 times. Note that these simulations have an illustrative purpose, to show how designs are affected by the components and biases. Effect sizes and biases definitions are not intended to be representative of true effects and are somewhat arbitrary. Additionally, by necessity we make certain as yet untested assumptions e.g. equal indirect genetic effects going between all siblings, equal strengths of population stratification and assortative mating among biological and adoptive parents. See below text under the heading 'Comparison of sibling, adoption, and non-transmitted allele designs in presence of simulated components and biases' and Supplementary Figure 9 for simulation results. Simulation code is fully available on Github.

<u>Sibling indirect genetic effects</u>
We simulate indirect genetic effects operating among three siblings in each family: individual 1, a biological sibling, and an unrelated adopted sibling. First, we create a matrix of sibling effects in which every effect is of the same magnitude (all siblings have an equal effect on each other regardless of adoption status, an implicit assumption), with zeros on the diagonal. To account for feedback effects (e.g. sibling 1 influences sibling 2, who influences sibling 1; this changes the coefficients of a variable on its own errors), we subtract the sibling effect matrix from an identity matrix and take its inverse. We then take the matrix product of the matrix with sibling

effects and the simulated sibling data to introduce the simulated mutual sibling effects into the data.

Assortative mating

Genetic assortative mating occurs when individuals with similar phenotypes mate more frequently than would be expected under a random mating scenario, and these phenotypes are heritable. In order to simulate assortment, we create phenotypes for the parents, rank the mothers and fathers by phenotype, and match couples according to rank. Since mating does not perfectly track with phenotypic rank, we add noise to the ranking of mothers and fathers prior to matching. Offspring genotypes are then simulated as random draws from the phenotypically matched couples' genotypes. Assortment is simulated to be of the same strength for the parents of adoptees as of non-adopted individuals, and we simulate random placement by un-ranking adoptees before matching them to adoptive families.

Population stratification

Population stratification can be conceptualized as systematic differences in allele frequencies between sub-populations. These frequency differences cause confounding in genetic studies when phenotypes also differ between sub-populations. We simulate such sub-populations in both the GWAS discovery and target PGS analyses samples. We first create genotypes in two groups, drawing upon two different sets of simulated minor allele frequency distributions. We also define these two groups as having different phenotypic means. We then run a single GWAS in these two populations. We create phenotypes and polygenic scores (based on the GWAS results) in a target sample of families, comprised of the same two sub-populations present in the GWAS. Our simulation allows for adoptees to be matched with adoptive parents both within- and between- sub-populations. We report simulation with adoptees matched with adoptive parents within the same sub-population.

#### Comparison of two implementations of the sibling design using simulation

In the sibling design presented by Selzam et al. (2019), indirect genetic effects are estimated by subtracting the within-sibling estimate from the between-sibling estimate (indexed using the average polygenic score for each sibling pair). However, the between-sibling effect is not necessarily the appropriate quantity to use (Carlin et al. 2005). An alternative is to subtract the within-sibling estimate from an estimate of the population effect obtained in a separate regression analysis using population data and ignoring family clustering. This approach was used in a recent within-sibling GWA study (Howe et al. 2021).

To ensure that we contrast our direct genetic effects with the appropriate quantity for accurate estimation of indirect genetic effects, we use simulated data to assess the use of the between-sibling effect and the population effect. Results are presented below in **Supplementary Figure 8**. From these simulations, it appears that contrasting the direct effects with the between-sibling effects leads to an overestimation of indirect parental genetic effects. Contrasting direct effects with population effects results in accurate estimation of indirect genetic effects. Consequently, we use this approach in our main analyses and simulations. Therefore, our model differs slightly from the Selzam et al. analyses.

Also notable is that, whilst the Selzam et al. article (and (Cheesman et al. 2020)) uses a different term – *passive gene-environment correlation* – the effect being estimated is a parental indirect genetic effect. Passive gene-environment correlation refers to how the genes that parents pass on to their children may also influence how they provide the rearing environment (Plomin et al. 1977).

Supplementary Figure 8. Estimation of the direct and parental indirect genetic effects in simulated data with different implementations of the sibling design.

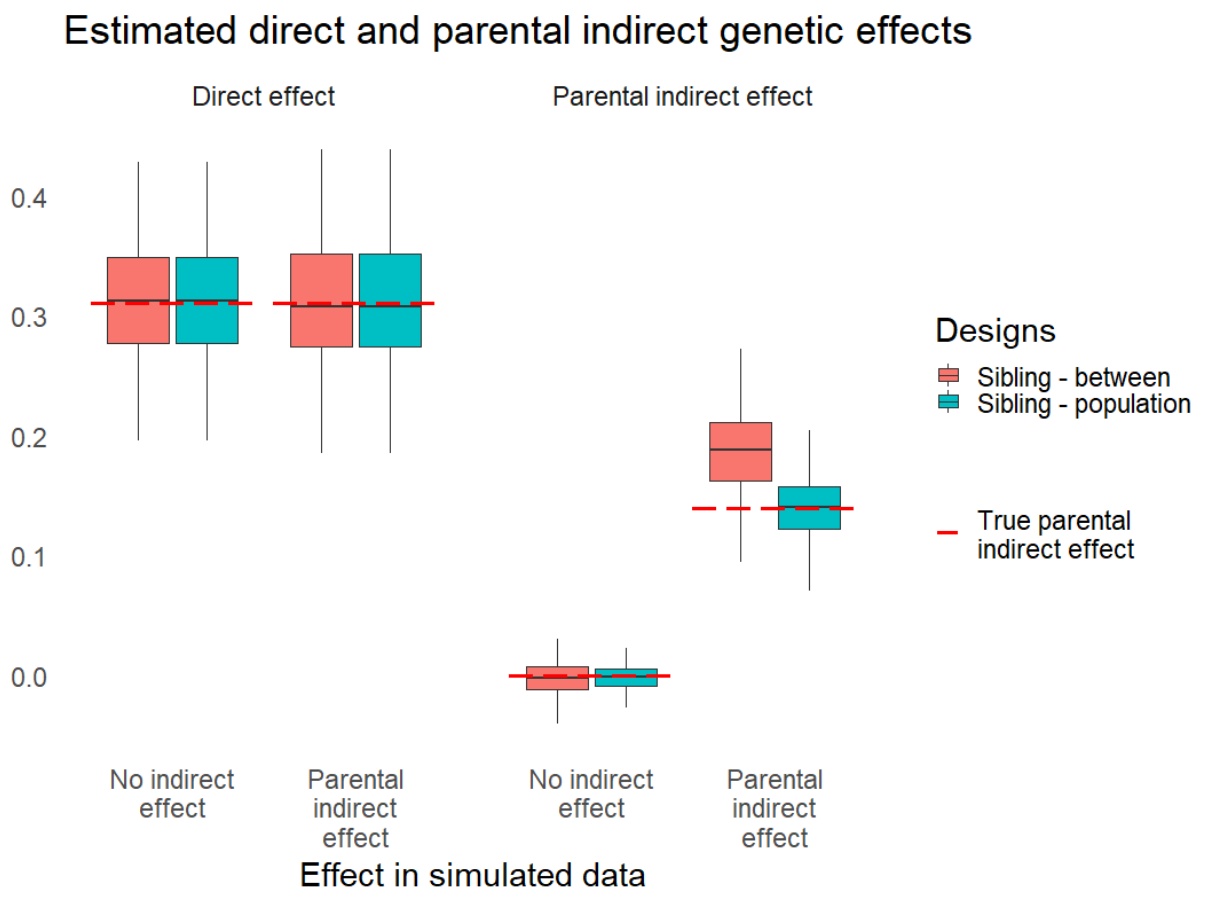

#### Comparison of sibling, adoption, and non-transmitted allele designs in presence of simulated components and biases

Using the simulated data, we compare the behaviour of the three designs used in our study to estimate direct and indirect genetic effects. For simplicity's sake our simulations consider one PGS (instead of both *Cog* and *NonCog* PGS). Additionally, we compare to a fourth design which we call "trios" in **Supplementary Figure 9**, in which the phenotype is simply regressed on child and parental PGS. As the simulation results prove, this simple approach gives identical estimates to the non-transmitted allele design, which also uses trios but requires prior identification of segments that are shared and non-shared between the generations.

**Supplementary Figure 9** (an extended version of **Figure 3** in the main text) displays the simulation results. The following text discusses the results, focusing on the main estimates of interest -- indirect genetic effects of parents.

<u>Prenatal parental indirect genetic effects</u>

We see prenatal effects as a component of interest, rather than as a bias, in estimates of indirect genetic effects. Nonetheless, for consistency with the rest of the simulations which do not consider prenatal indirect genetic effects, the red dashed line in **Supplementary Figure 9** indicates the true postnatal effect only. Simulation results show that the sibling- and trio-based designs capture indirect genetic effects occurring in both prenatal and postnatal periods. In contrast, the adoption design only captures postnatal indirect genetic effects. This is because, for both adoptees and non-adopted individuals, the prenatal environment is provided by the biological mother, so estimated polygenic score-phenotype associations for both adoptees and non-adopted individuals contain prenatal maternal indirect genetic effects. Consequently, computing the indirect genetic effect as the population effect of the polygenic score (β in non-adopted individuals) minus the direct genetic effect (β for adoptees) means that prenatal effects are cancelled out. This result suggests that prenatal indirect genetic effects could partially explain lower estimates of indirect genetic effects from the adoption design compared to the other designs.

<u>Sibling indirect genetic effects</u>

We find that positive sibling effects result in upwardly biased estimates of indirect parental genetic effects. This bias is considerably larger for the sibling design than the adoption and trio designs. Bias in the sibling design is likely to be because positive sibling effects increase the similarity of siblings, reducing the effect of within-sibling polygenic differences (Boardman and Fletcher 2015; Kohler et al. 2011). Bias in the trio design likely arises because non-transmitted alleles are not only shared with parents, but also partially with (full) siblings, such that $\beta NT$ might capture sibling as well as parental indirect genetic effects. It is interesting that sibling effects inflate adoption-based estimates despite the fact that adoptees are not genetically related to their siblings. These simulation results lead to the notion that higher estimates of parental indirect genetic effects in the sibling than adoption and trio designs is evidence of sibling indirect genetic effects. In our empirical data, we do not find differences between sibling- and trio-based estimates of indirect parental genetic effects. Along with our sensitivity analyses, this suggests an absence of sibling genetic effects on educational outcomes in our datasets.

Assortative mating

We found that the bias from assortative mating in the indirect genetic effect estimate was lower in the adoption design than in the non-transmitted allele and sibling designs. Interestingly, the bias in the adoption design from assortative mating is zero in the absence of a parental indirect genetic effect, but slightly above zero when a parental indirect genetic effect was also specified. In other words, the presence of parental indirect genetic effects is required for assortative mating to bias estimates from the adoption design. We simulated the same strength of assortative mating for the parents of both adopted and non-adopted individuals, so the result cannot be due to elevated assortment in the latter group (leading to residual assortment in the indirect effect estimate when calculating β*non-adopted* - β*adopted*). Such differences could exist in the real data, but there is scarce and inconsistent evidence regarding assortment in biological parents of adoptees versus other parents (Plomin et al. 1977; Ho et al. 1979). Overall, the results suggest that assortative mating could explain lower estimates of indirect genetic effects from the adoption design compared to the other designs.

Population stratification

Simulation results show that estimates of parental indirect genetic effects based on the adoption design capture less bias from population stratification than sibling- and trio-based designs. In the sibling design, the parental indirect genetic effect is estimated as the population effect minus the direct within-family effect of the polygenic score. This means that the indirect genetic effect is likely to be inflated by population stratification, as this is captured in the population effect but not the within-family effect. Also, the effects of the non-transmitted allele PGS are influenced by population stratification, so the indirect genetic effect estimate is inflated. In contrast, population stratification only biases indirect genetic effect estimates from the adoption design to a small extent. Assuming that population stratification is similar in adoptees and non-adopted individuals, its effect will cancel out when estimating the indirect genetic effect as β*non-adopted* - β*adopted*. The assumption of equal population stratification and assortative mating bias in adopted and non-adopted groups cannot be tested due to the lack of parental data in UKB, but is bolstered by the simulation results, and by the fact that both adoptees and non-adopted individuals are from British ancestry. Our simulation results suggest that population stratification partly explains the lower estimates of indirect genetic effects from the adoption design compared to the other designs in our empirical study.

#### Supplementary Figure 9. Simulation results: Comparison of sibling, adoption, and non-transmitted allele designs in presence of components and biases

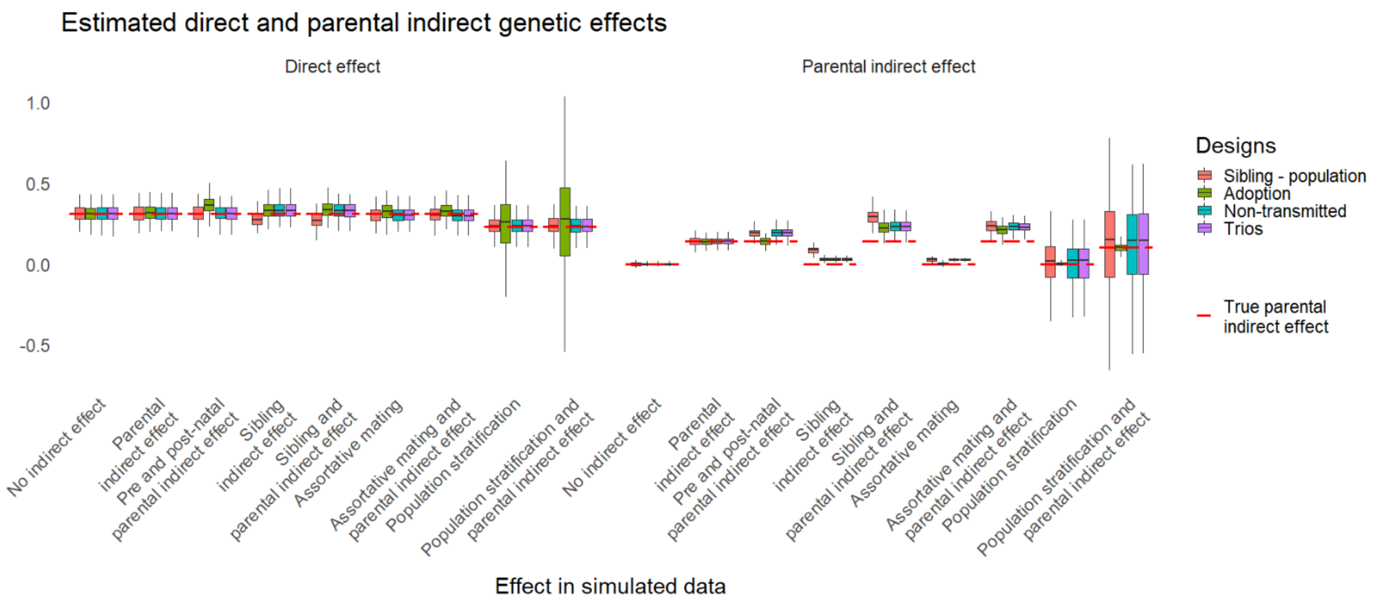
